# Metabolic and signalling network map integration: application to cross-talk studies and omics data analysis in cancer

**DOI:** 10.1101/274902

**Authors:** Nicolas Sompairac, Jennifer Modamio, Emmanuel Barillot, Ronan M. T. Fleming, Andrei Zinovyev, Inna Kuperstein

**Affiliations:** Institut Curie, 26 rue d’Ulm, F-75005 Paris, France, Inserm, U900, F-75005, Paris France, Mines Paris Tech, F-77305 cedex Fontainebleau, France, PSL Research University, F-75005 Paris, France; Centre for Systems Biomedicine, University of Luxembourg, 6 avenue du Swing, L-4367 Belvaux, Luxembourg

**Keywords:** Keywords Signalling, Metabolism, Networks, Comprehensive map, Systems Biology, Cancer, Multi-level omics data, Data visualisation

## Abstract

**Background:** The interplay between metabolic processes and signalling pathways remains poorly understood. Global, detailed and comprehensive reconstructions of human metabolism and signalling pathways exist in the form of molecular maps, but they have never been integrated together. We aim at filling in this gap by creating an integrated resource of both signalling and metabolic pathways allowing a visual exploration of multi-level omics data and study of cross-regulatory circuits between these processes in health and in disease.

**Results:** We combined two comprehensive manually curated network maps. Atlas of Cancer Signalling Network (ACSN), containing mechanisms frequently implicated in cancer; and ReconMap 2.0, a comprehensive reconstruction of human metabolic network. We linked ACSN and ReconMap 2.0 maps via common players and represented the two maps as interconnected layers using the NaviCell platform for maps exploration. In addition, proteins catalysing metabolic reactions in ReconMap 2.0 were not previously visually represented on the map canvas. This precluded visualisation of omics data in the context of ReconMap 2.0. We suggested a solution for displaying protein nodes on the ReconMap 2.0 map in the vicinity of the corresponding reaction or process nodes. This permits multi-omics data visualisation in the context of both map layers. Exploration and shuttling between the two map layers is possible using Google Maps-like features of NaviCell. The integrated ACSN-ReconMap 2.0 resource is accessible online and allows data visualisation through various modes such as markers, heat maps, bar-plots, glyphs and map staining. The integrated resource was applied for comparison of immunoreactive and proliferative ovarian cancer subtypes using transcriptomic, copy number and mutation multi-omics data. A certain number of metabolic and signalling processes specifically deregulated in each of the ovarian cancer sub-types were identified.

**Conclusions:** As knowledge evolves and new omics data becomes more heterogeneous, gathering together existing domains of biology under common platforms is essential. We believe that an integrated ACSN-ReconMap 2.0 resource will help in understanding various disease mechanisms and discovery of new interactions at the intersection of cell signalling and metabolism. In addition, the successful integration of metabolic and signalling networks allows broader systems biology approach application for data interpretation and retrieval of intervention points to tackle simultaneously the key players coordinating signalling and metabolism in human diseases.

## Background

There is still a gap in understanding the coordination between metabolic functions and signalling pathways in mammalian cells. Metabolic processes and cell signalling pathways contain a large number of molecular species together with their complex relationships. No single mind can accurately account for all these molecular interactions whilst drawing conclusions from a process of descriptive thought. To tackle the complexity of these multi-molecular interactions networks, a systems biology approach is needed. In addition, there is a high number of omics data such as transcriptome, proteome, metabolome, etc. accumulated for many human diseases as age-related disorders (e.g. neurodegeneration or cancer). Modelling and interpretation of these data combining metabolic and signalling networks together can help to decipher the mechanisms responsible for deregulations in human disorders by considering a broader range of molecular processes types.

Much of produced high-throughput molecular data in many medical and biological applications remain under-explored due to the lack of insightful methods for data representation in the context of formally represented biological knowledge. Carefully designed maps of complex molecular mechanisms such as the whole-cell reconstructions of human metabolism in ReconMap 2.0 [1][2] or the global reconstruction of cell signalling of cancer in ACSN [3] potentially provide ways to better exploit existing and new multi-omics data, by overlaying it on top of large molecular maps.

ACSN is a resource and a web-based environment that contains a collection of interconnected signalling network maps (https://acsn.curie.fr). Cell signalling mechanisms are depicted on the maps at the level of biochemical interactions, forming a large network of 4600 reactions covering 1821 proteins and 564 genes and connecting several major cellular processes [3]. ACSN is composed of 5 interconnected maps of major biological processes implicated in cancer. The maps are further divided into functional modules that represent signalling pathways collectively responsible for the execution of a particular process. In total, there are 52 functional modules in the ACSN resource (See Table 1 for terms definition). Each of these modules can be visualised in the context of the global ACSN map or accessed as individual maps. The Atlas is a “geographic-like” interactive “world map” of molecular interactions. ACSN is supported by NaviCell platform for easy map navigation and its annotations using Google maps^™^ engine. The logic of navigation as scrolling and zooming; features as markers, pop-up bubbles and zoom bar are adapted from the Google map. Finally, NaviCell includes a powerful module for data visualisation. Users can map and visualise different types of “omics” data on the NaviCell maps [4][5].

The manually curated genome-scale reconstruction Recon2.04 is a representation of the human metabolism. It accounts for 1,733 enzyme-encoding genes associated to 7,440 reactions which are distributed in 100 subsystems, referring to metabolic pathways. Furthermore, Recon2.04 accounts with 2,626 unique metabolites distributed over eight cellular compartments [2]. Subsequently, to visualise the resource, a comprehensive metabolic map termed ReconMap 2.0 was generated from the Recon2.04 resource [1]. In the ReconMap 2.0 reactions (hyper-edges) were manually laid out using the biochemical network editor CellDesigner [6]. ReconMap 2.0 is currently distributed in a Systems Biology Graphical Notation compliant format and its content is also accessible via a web interface (https://vmh.uni.lu/#reconmap). All major human metabolic pathways are considered and represented as a seamless network where different pathways are interconnected via common molecules. There are 96 subsystems on the ReconMap 2.0, each of them representing a specific metabolic pathway (See Table 1 for terms definition).

By integrating these resources together, it will be possible to elucidate the crosstalk between metabolic and signalling networks. In addition, the integrated networks, provided in a common graphical language and available in standard exchange formats, makes them accessible for multiple systems biology tools. It opens an opportunity to model coordination between signalling pathways and metabolism using various systems biology approaches. Among others, there are several methods for multi-level omics data analysis in the context of the biological network maps that allow defining “hot” areas in molecular mechanisms and point to key regulators in physiological or in pathological situations [7][8][9] and beyond.

## Results

### 1. General workflow for integration of ACSN and ReconMap 2.0 networks

With the aim to integrate signalling and metabolic networks there is a need to find common players (proteins) that participate in the regulation of metabolic processes and simultaneously involved in signal transduction pathways. Thus, the networks can be interconnected via these common players. In addition, some solution for visualisation of proteins participating in the catalytic process in ReconMap 2.0 should be provided, since there is no such representation up to date.

The rationale behind the proposed methodology is to take an advantage of the CellDesigner SBML format for networks representation and develop a robust automated algorithm for an efficient finding of coordinates for new entities avoiding an overlap with existing elements and visualising these entities in the vicinity of the corresponding reactions they regulate. The integrated networks can be provided as interconnected layers supported by NaviCell platform for navigation and data integration.

The suggested methodology is applied for ACSN and ReconMap 2.0 resources integration. However, this is a generic method applicable for integration of different types of networks prepared in CellDesigner SBML format. In the following sections of the paper, we explain the challenges and describe how each step mentioned in the workflow was addressed.

The workflow in the Section 2 includes the following major steps (see Table 1 for terms definition):

- Identification of common proteins between ACSN and ReconMap 2.0 networks
- Finding metabolic and molecular processes crosstalk between ACSN and ReconMap 2.0
- Displaying proteins nodes on the ReconMap 2.0 map
- ACSN-ReconMap 2.0 networks integration and visualisation using NaviCell

**Table 1.** Terms definitions used in the paper

| Graphical standards and exchange formats |  |
| --- | --- |
| XML format | Markup language that defines a set of rules for encoding documents used to store and transport data by describing the content in terms of what data is being described. |
| SBGN | Systems Biology Graphical Notation (SBGN) is a standard graphical syntax for representation of biological processes and interactions. SBGN is compatible with multiple pathway drawing and analytical tools, http://sbgn.github.io/sbgn/ |
| SBML | Systems Biology Markup Language (SBML) is a representation format, based on XML, for communicating and storing computational models of biological processes. It is a free and open standard language with widespread software support, http://sbml.org |
| Standard identifier (ID) | Community-accepted nomenclature for scientific naming of biomolecules as genes, proteins, chemicals, drugs etc. The sources for standard IDs are repositories as UNIPROT, CHEB, HUGO, http://identifiers.org |
| Data and models exchange formats | Standard formats for data and models to facilitate networks and software interoperability. There are two major standard networks exchange formats, BIOPAX for complex networks and SIF for simple binary interactions. The CellDesigner xml format is commonly-used exchange format compatible with multiple network analysis tools. |
| Signalling and metabolic network map |  |
| Map | Diagram of detailed molecular interactions with meaningful layout reflecting a certain biological process, which is graphically represented in CellDesigner tool. |
| Map module (in ACSN) | Part of the map representing a sequence of molecular interactions responsible for execution of a particular function. |
| Metabolic pathway Subsystem (in ReconMap 2.0) | Set of reactions forming a metabolic function.<br>Set of metabolic reactions associated to (representing) a specific metabolic pathway. |
| Map node | Graphical representation of a molecule on the map. |
| Map entity and alias | Unique representation of a molecule on the map. As each molecule can be present multiple times on a map, each individual representation is called an alias of the entity. This definition corresponds to the CellDesigner feature. |
| Networks merging procedure |  |
| Voronoi cell | Individual shape allocated to a seed from the Voronoi method. Each cell is a space that contains only the seed and can be used to generate points inside without overlapping with close seeds. |
| Voronoi tessellation | A partitioning of a plane into regions based on distance to points in a specific subset of the plane. That set of points (called seeds) is specified beforehand, and for each seed there is a corresponding region consisting of all points closer to that seed than to any other. These regions are called Voronoi cells. In our case, each seed is a molecule or a reaction's central glyph. |
| Centroid | Barycenter of a cluster. |
| Merging function of BiNoM | Function allowing to take two or more CellDesigner maps and merge them in one unique map. This function modifies each entities' id and alias but keeps the name, coordinates and notes. |
| NaviCell |  |
| Semantic zoom | A mechanism providing several map views with different levels of details depiction achieved by gradual exclusion of details while zooming out. It simplifies navigation through large maps of molecular interactions by providing several levels of details, resembling navigation through geographical maps. Exploring the map from a detailed toward a top-level view is achieved by gradual exclusion and modification (simplification and abstraction) of details. One of the main principle of semantic zooming is in that every detail which is shown on the map at a current zoom level, should be readable. |
| Marker | Symbol indicating location of chosen objects on the map; adapted from Google maps. |
| Pop-up bubble | Small window that opens by clicking on marker. Contains short description and hyperlinks related to the marked entity. |
| Annotation post | Detailed map entity annotation created in CellDesigner by map manager. The annotation is converted to annotation post and displayed in the associated blog by NaviCell. |

## 2. Step-by-step procedure for network integration

### 2.1 Identification of common proteins between ACSN and ReconMap 2.0 networks

ACSN and ReconMap 2.0 maps contain information on proteins implicated in the regulation of reactions. First, the systematic use of the common identifiers as standard protein names (HUGO) for all proteins in both resources was verified and inconsistencies corrected. Thus, the proteins found in both resources ACSN and ReconMap 2.0, were compared, quantified and visualised. We detected 252 proteins in common between the two networks (Additional file 1).

### 2.2 Finding metabolic and molecular processes crosstalk between ACSN and ReconMap 2.0

ACSN and ReconMap 2.0 networks have a particular hierarchical structure. ACSN is divided into functional modules, whereas ReconMap 2.0 is divided into subsystems. Each of these structures are a subset of processes from the global network, implicated in regulation and execution of a specific molecular or metabolic pathway respectively (See Table 1 for terms definition). To address the question which metabolic processes are connected to which signalling mechanisms, the enrichment analysis of ACSN modules and Recon 2.0 subsystems was performed using the 252 common proteins (Additional file 1). The composition of ACSN modules and ReconMap 2.0 subsystems are provided as gene sets in Additional files 2 and 3 in gmt format and the enrichment was calculated using a Hypergeometric Test. The analysis demonstrated that proteins shared between the two maps are implicated in 22 modules of ACSN and in 51 subsystems of ReconMap 2.0 (Figure 1 and Table 2). Majority of proteins on both resources are participating in catalysis of biochemical or metabolic reactions. The information for the protein-reaction association is encoded in the network structure and in the annotations on the CellDesigner XML files. Information for reactions in each ACSN module and ReconMap2.0 subsystem was retrieved and quantified. The number of reactions in ReconMap 2.0 subsystems regulated by proteins from ACSN modules is shown in Additional file 4.

**Figure 1.**
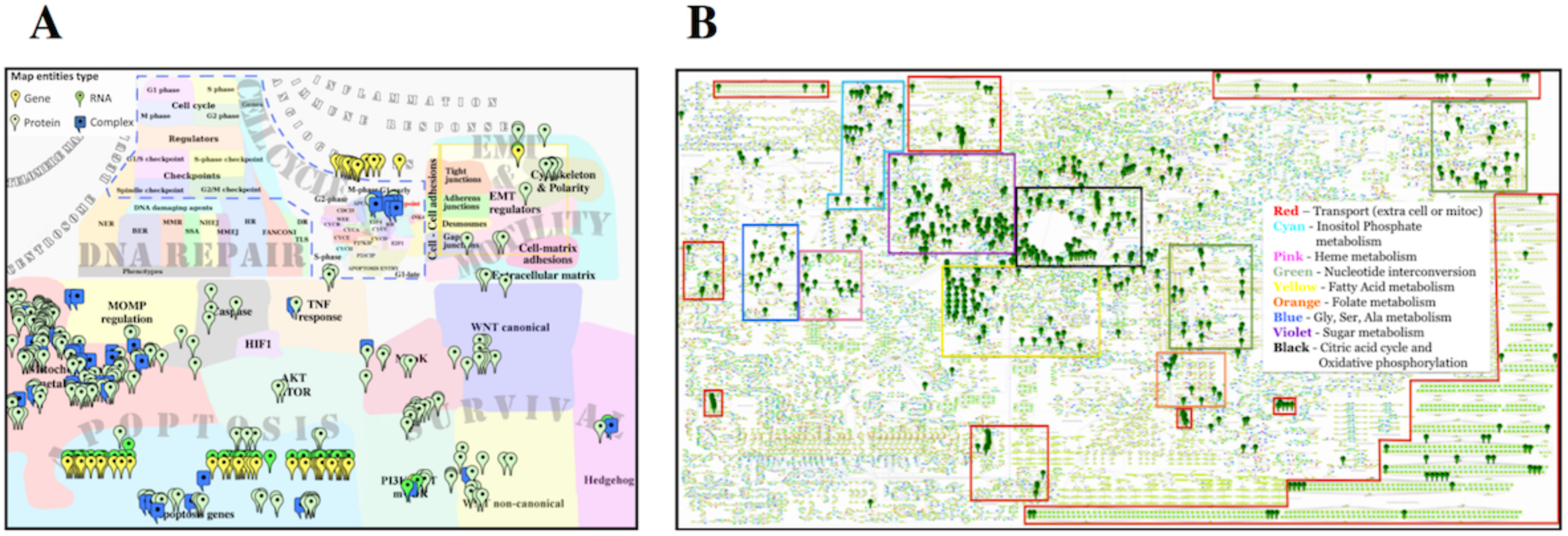
Distribution of proteins common between ACSN and ReconMap 2.0 networks. Proteins are found in various modules of ACSN (A) and metabolic pathways of ReconMap 2.0 (B). Markers indicate the proteins (enzymes catalysing metabolic reactions in ReconMap 2.0) that also found in the signalling pathways of ACSN.

Since ACSN is the resource focused on processes implicated in cancer; as expected, the ACSN modules containing mechanisms related to Mitochondria metabolism and Apoptosis have many shared proteins with ReconMap 2.0. Interestingly, ACSN modules related to cell survival processes as Wnt-Non-canonical pathway and modules related to the invasion and mobility processes as EMT regulators are also enriched by proteins regulating metabolic pathways of ReconMap 2.0. On the ReconMap 2.0, the shared proteins are enriched in energy-providing processes as Citric Acid Cycle (Krebs cycle) and Oxidative Phosphorylation and in processes related to Pentose Phosphate Pathway, Fructose and Mannose metabolism, Glycolysis and Gluconeogenesis. Finally, the subsystem Inositol Phosphate Metabolism is enriched by the shared proteins (Figure 1 and Table 2).

The same trend is observed at the level of the reactions regulation, finding proteins from Apoptosis, Cell cycle and Mitochondrial processes implicated in catalysis of a large number of reactions in the ReconMap 2.0 is expected as these pathways are known to be closely related to the metabolism. However, it is interesting to note that some less intuitive connections between ReconMap 2.0 subsystems and ACSN modules were retrieved. For example, proteins implicated in cell survival modules of ACSN as Hedgehog, MapK, PI3K-AKT-mTOR and WNT regulate reactions in the Inositol phosphate metabolism subsystem from ReconMap 2.0. In addition, the proteins from cell migration-related and epithelial-to-mesenchymal transition (EMT)-related processes we involved in the regulation on the reactions in five different phospholipids and amino acid metabolic pathways, indicating most probably that the invasion process requires very active metabolism in migrating cancer cells. These connections are less obvious and may help to highlight interesting connections between signalling and metabolic processes in cancer (Additional file 4).

By extracting information about crosstalk between ACSN modules and ReconMap 2.0 subsystems, it was possible to generate a network where nodes represent ACSN modules and ReconMap 2.0 subsystems connected by edges if they shared common proteins (Figure 2, Table 2 and Additional File 5). The obtained network contains one large connected component and also a number of modules and subsystems that are not connected to each other.

**Figure 2.**
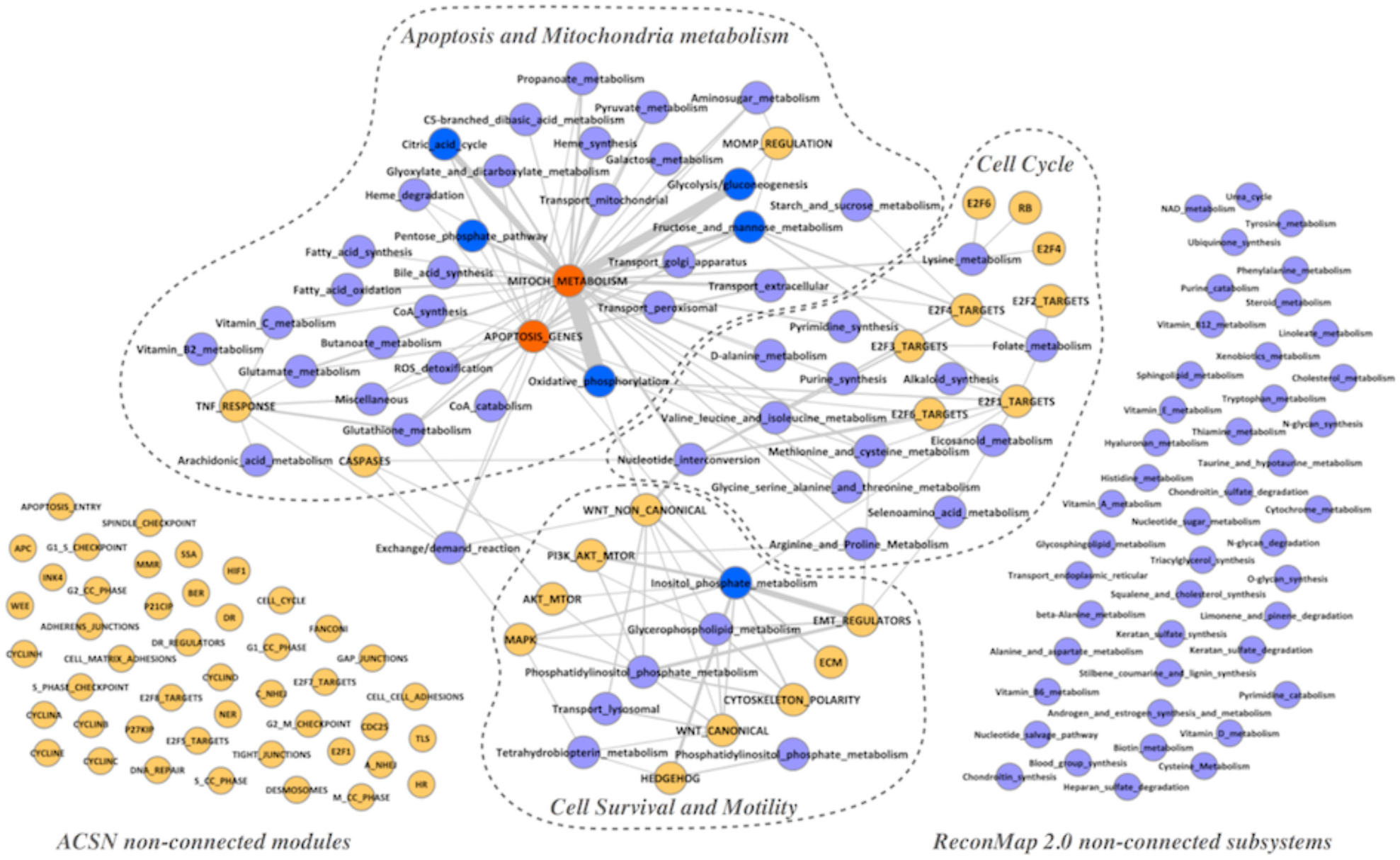
Network of crosstalk between ACSN modules and ReconMap 2.0 subsystems. ACSN modules and ReconMap 2.0 subsystems are represented as the nodes of the networks and connected by edges if there are shared proteins between them. Edges width is proportional to the number of proteins in the intersection. Nodes representing ACSN modules are coloured in Orange and ReconMap 2.0 subsystems are coloured in Light Blue. The nodes representing enriched ACSN modules are coloured in Red and enriched ReconMap 2.0 subsystems are coloured in Dark Blue

The interconnections between many signalling and metabolic processes on the network form communities. It was possible to identify three major communities each containing ACSN modules interconnected with ReconMap 2.0 subsystems, that we called ‘Apoptosis and Mitochondrial Metabolism’, ‘Cell Cycle’ and ‘Cell Survival and Motility’ according to their biological functions (Figure 2). The nodes Mitochondria Metabolism and Apoptosis Genes, two modules of ACSN, are shown to be enriched with common proteins and connected to 36 and 24 subsystems of ReconMap 2.0 respectively. A total of 20 metabolic subsystems were found grouped forming a big community related to ‘Apoptosis and Mitochondrial Metabolism’ This result is not surprising since within the mitochondrion we find main cellular pathways such as Citric acid cycle, Oxidative phosphorylation and Fatty acid oxidation. Furthermore, mitochondrion is a key organelle regulating cell death via two ways. On one hand, the lack of ATP which is mainly produced in the mitochondria via Oxidative phosphorylation will lead to cell death through necrosis [10] and on the other hand, a variety of signalling apoptotic processes are linked to mitochondria [11], such as for example the family Bcl-2 which regulates apoptosis through mitochondrial permeability [10][12]. The Inositol Phosphate Metabolism subsystem is linked to 11 modules of ACSN, most of them being part of the ‘Cell Survival and Motility’ community. In accordance with this result, Inositol Phosphate functions act as second messengers for a variety of extracellular signals. Their effect in cell motility is due to the interaction of cellular membranes with proteins of the cytoskeleton [13]. Furthermore, members of the inositol phosphate metabolism pathway regulate the phosphatidylinositol-3-kinase (PI3K)/Akt signalling pathway, therefore cellular quiescence, proliferation, cancer, and longevity processes [14]. Interestingly enough, the subsystems responsible for nucleotides synthesis and metabolism related to the ‘Cell Cycle’ community are actually crosstalking with all three communities. Amino acids (aa) are not only used as a main energy source via oxidation and integration within the citric acid cycle in the mitochondrion; but also, they play a key role in several signalling pathways. Amino acids deprivation leads to cellular death through apoptosis and autophagy [15]. Moreover, their relation with cell motility has been also previously reported [16]. In addition, their association with cell proliferation seems to be correlated with the differential synthesis of proteins at different stages of the cell cycle [17][18]. This observation demonstrates the central role of nucleotide metabolism in the major cell processes.

The list of signalling modules and metabolic sub-systems that do not intersect in the current versions of both resources suggests performing literature mining with the aim to figure out whether a potential coordination between these processes is documented.

**Table 2.** Crosstalk between ACSN modules and ReconMap 2.0 metabolic pathways. Using the composition of each module and subsystem (Additional files 2 and 3) and the list of the 252 shared proteins between both maps, modules and subsystems having common proteins were linked together. Thus, a crosstalk between both maps has been uncovered with the number of shared proteins showed here. *Modules and subsystems in bold are the result of an Enrichment Analysis of the 252 common proteins between ACSN and ReconMap 2 using the Survival Function Method. The p-value is obtained through a standard Hypergeometric test on modules/subsystems containing at least 10 proteins (p-value inferior to 1e-2).

| ACSN Modules | Number of shared proteins in ACSN modules | ReconMap 2.0 subsystem | Number of shared proteins in ReconMap 2.0 subsystems |
| --- | --- | --- | --- |
| MITOCH_METABOLISM* | 186 | Propanoate_metabolism;Transport_golgi_apparatus;Valine_leucine_and_isoleucine_metabolism;Glutamate_metabolism;C5-branched_dibasic_acid_metabolism;Fatty_acid_synthesis;Citric_acid_cycle*;Vitamin_C_metabolism;Nucleotide_interconversion;Glycine_serine_alanine_and_threonine_metabolism;Galactose_metabolism;Pentose_phosphate_pathway*;Miscellaneous;Exchange/demand_reaction;Aminosugar_metabolism;Glyoxylate_and_dicarboxylate_metabolism;Lysine_metabolism;Transport_extracellular;Fatty_acid_oxidation;Glutathione_metabolism;Fructose_and_mannose_metabolism*;Transport_mitochondrial;Pyrimidine_synthesis;Heme_synthesis;Butanoate_metabolism;Pyruvate_metabolism;D;alanine_metabolism;Glycolysis/gluconeogenesis*;Transport_peroxisomal;Oxidative_phosphorylation*;ROS_detoxification;Bile_acid_synthesis;Methionine_and_cysteine_metabolism;Purine_synthesis;Arginine_and_Proline_Metabolism;Heme_degradation | 245 |
| APOPTOSIS_GENES* | 46 | Transport_golgi_apparatus;Glutamate_metabolism;CoA_catabolism;Citric_acid_cycle*;Glyoxylate_and_dicarboxylate_metabolism;Glycine_serine_alanine_and_threonine_metabolism;Inositol_phosphate_metabolism*;Pentose_phosphate_pathway*;Aminosugar_metabolism;Miscellaneous;Transport_extracellular;Propanoate_metabolism;Fructose_and_mannose_metabolism*;Heme_synthesis;Exchange/demand_reaction;Pyruvate_metabolism;Glycolysis/gluconeogenesis*;ROS_detoxification;Glutathione_metabolism;Methionine_and_cysteine_metabolism;Oxidative_phosphorylation*;CoA_synthesis;Heme_degradation | 56 |
| WNT_NON_CANONICAL | 23 | Transport_lyosomal;Exchange/demand_reaction;Phosphatidylinositol_phosphate_metabolism;Glycerophospholipid_metabolism;Nucleotide_interconversion;Inositol_phosphate_metabolism* | 27 |
| EMT_REGULATORS | 22 | Glycerophospholipid_metabolism;Inositol_phosphate_metabolism*;Phosphatidylinositol_phosphate_metabolism;Selenoamino_acid_metabolism;Methionine_and_cysteine_metabolism | 36 |
| CYTOSKELETON_POLARITY | 5 | Glycerophospholipid_metabolism;Inositol_phosphate_metabolism*;Phosphatidylinositol_phosphate_metabolism* | 12 |
| E2F4_TARGETS | 11 | Starch_and_sucrose_metabolism;Fructose_and_mannose_metabolism*;Folate_metabolism;Nucleotide_interconversion;Transport_extracellular | 12 |
| E2F1_TARGETS | 14 | Fructose_and_mannose_metabolism*;Folate_metabolism;Eicosanoid_metabolism;Nucleotide_interconversion;Selenoamino_acid_metabolism;Methionine_and_cysteine_metabolism;Inositol_phosphate_metabolism*;Oxidative_phosphorylation*;Alkaloid_synthesis | 15 |
| AKT_MTOR | 3 | Inositol_phosphate_metabolism*;Phosphatidylinositol_phosphate_metabolism;Glutathione_metabolism | 4 |
| PI3K_AKT_MTOR | 10 | Phosphatidylinositol_phosphate_metabolism;Inositol_phosphate_metabolism*;Arginine_and_Proline_Metabolism | 13 |
| CASPASES | 3 | Glycerophospholipid_metabolism;Exchange/demand_reaction;Nucleotide_interconversion | 3 |
| ECM | 5 | Inositol_phosphate_metabolism* | 5 |
| TNF_RESPONSE | 13 | Vitamin_B2_metabolism;Glutamate_metabolism;Miscellaneous;Glutathione_metabolism;Arachidonic_acid_metabolism;Exchange/demand_reaction;Vitamin_C_metabolism | 15 |
| E2F3_TARGETS | 5 | Folate_metabolism;Nucleotide_interconversion | 5 |
| MAPK | 8 | Tetrahydrobiopterin_metabolism;Inositol_phosphate_metabolism*;Phosphatidylinositol_phosphate_metabolism;Glycerophospholipid_metabolism;Exchange/demand_reaction | 11 |
| MOMP_REGULATION | 1 | Fructose_and_mannose_metabolism*;Aminosugar_metabolism;Glycolysis/gluconeogenesis* | 3 |
| WNT_CANONICAL | 8 | Transport_lyosomal;Phosphatidylinositol_phosphate_metabolism;Glycerophospholipid_metabolism;Tetrahydrobiopterin_metabolism;Inositol_phosphate_metabolism*;Oxidative_phosphorylation* | 14 |
| HEDGEHOG | 10 | Tetrahydrobiopterin_metabolism;Inositol_phosphate_metabolism*;Phosphatidylinositol_phosphate_metabolism | 13 |
| E2F6_TARGETS | 5 | Oxidative_phosphorylation*;Nucleotide_interconversion | 5 |
| RB | 1 | Lysine_metabolism | 1 |
| E2F6 | 2 | Lysine_metabolism | 2 |
| E2F2_TARGETS | 1 | Folate_metabolism | 1 |
| E2F4 | 1 | Lysine_metabolism | 1 |

### 2.3 Displaying proteins nodes on the ReconMap 2.0 map

ACSN and ReconMap 2.0 are both used as visual objects for exploration of processes as well as for data integration and visualisation in the context of the maps. After identification of the cross-talks between the two resources, it is important to ensure that all components of the maps are represented in a visual manner suitable for meaningful visualisation of omics data.

Due to the different nature of the networks, protein nodes are explicitly visualised on the ACSN map. However, in the ReconMap 2.0 the Standard Names (Identifiers) of proteins regulating metabolic reactions are included into the reaction annotations, but not represented visually on the map canvas. This precludes visualisation of omics data in the context of ReconMap 2.0 map. We developed a procedure for displaying the protein nodes on the ReconMap 2.0 map in the vicinity of the corresponding reaction edges, that now permits a multi-omics data visualisation in the context of both ACSN and ReconMap 2.0 layers.

#### Extraction of information regarding reactions and implicated genes in the metabolic network

##### -Recuperation of the information from the Recon2.04 model

ReconMap 2.0 is the graphical representation of the literature-based genome-scale metabolic reconstruction Recon2.04, which is freely available at (https://vmh.uni.lu/#downloadview). It is stored as a MatLab “.mat” file which contains a direct link between metabolic reactions and gene Entrez, specified by gene-rules. Therefore, it is possible to generate a direct protein-reaction association based on the gene codifying for the protein. As ACSN uses HUGO Standard Identifiers, Entrez IDs in ReconMap 2.0 were first converted to HUGO.

It is important to stress that this approach is based on a simplified assumption that if a protein is associated to a metabolic reaction in ReconMap 2.0, it may have a role in catalysis of the reactions. However, it is clear that the biological regulation is much more sophisticated than this basic assumption. For example, there are many protein complexes collectively regulating propagation of metabolic reaction and only part of them are actual enzymes that execute the catalysis, whereas others are co-factors of regulatory sub-units. Moreover, the activation states of proteins that is often regulated by post-translational modifications are also not taken into account in this simplified approach.

##### -Recuperation of entities positions in ReconMap 2.0 from the XML network file

In the graphical representation of reactions in CellDesigner, each reaction contains a central glyph in the form of a square. This glyph is normally used to allocate the position of the markers (see Table 1 for terms definition). However, its location is not explicitly saved in the network XML file. A specific function of NaviCell factory can calculate the coordinates of these glyphs and extract them in a separated file. These coordinates can be later used as a reference positions to assign proteins nodes position in the ReconMap 2.0 map canvas.

#### Automated calculation of proteins coordinates in vicinity of corresponding reactions at ReconMap 2.0network

##### -Computing Voronoi cells for all elements

By using the Voronoi method, each element of the network (molecules, reaction glyphs, etc.) is associated to a Voronoi cell. By using this method, it could be guaranteed to avoid any overlapping with already existing entities in the network when adding new proteins (Figure 3).

##### -Creation of randomly distributed points inside each reaction’s Voronoi cell

When each entity has a cell assigned, cells of reactions’ central glyphs are utilised. Each cell has a certain number of points assigned randomly inside the cell. For our purpose, 100 points were deemed sufficient (Figure 3).

##### -Application of K-means algorithm to create K clusters

Each reaction has a certain number of proteins implicated in its catalysis. Using the information from the model, the K-means algorithm was applied to identify the number of clusters centres corresponding to the number of protein nodes (Figure 3).

##### -Assigning protein positions using centroids coordinates of each cluster

After the protein clusters are found, their centroids (see Table 1 for terms definition) are calculated and will be saved as the coordinated of the proteins tied to the specific reaction as catalysts (Figure 3).

**Figure 3.**
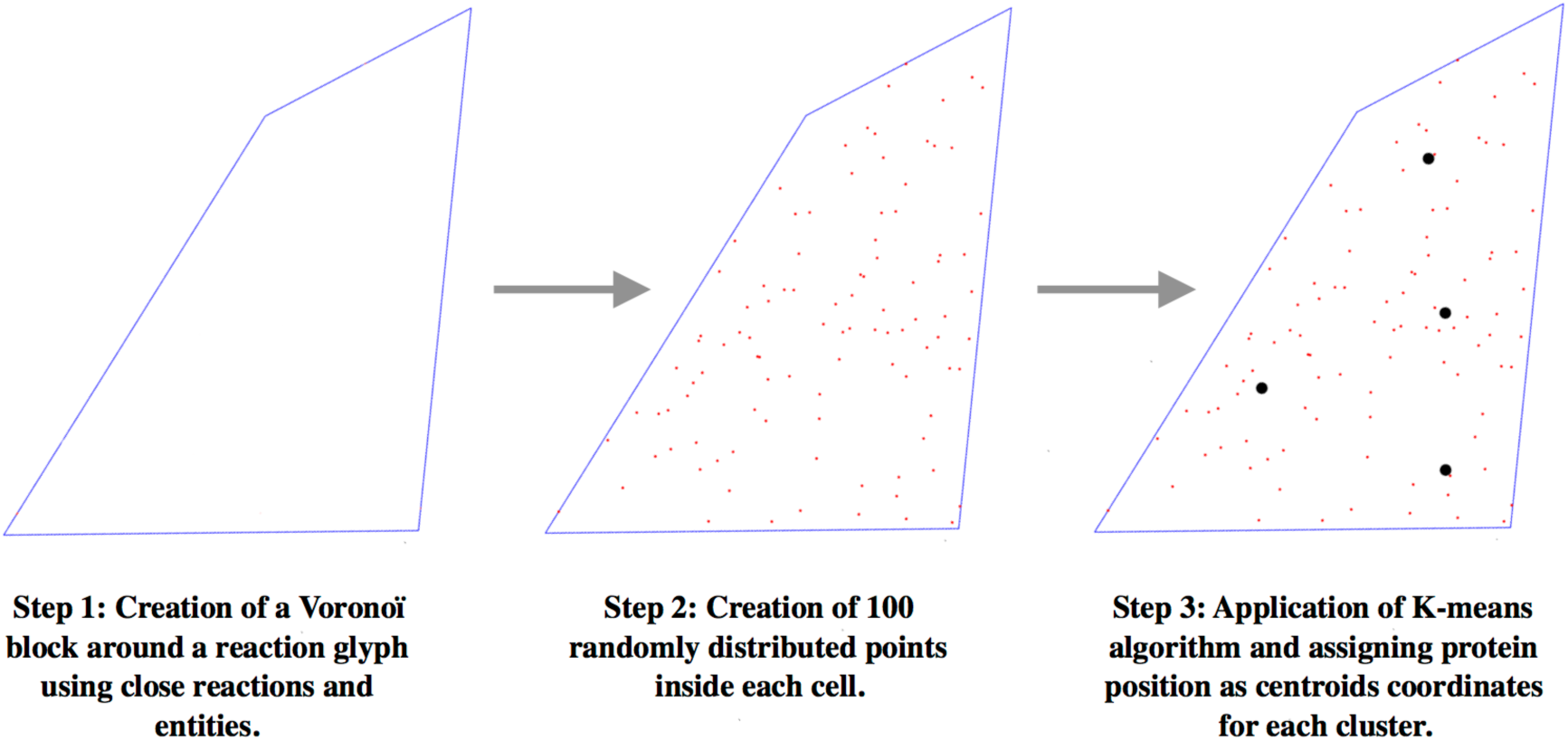
Illustration of the three steps for automated proteins addition in the vicinity of a reaction. The first step is to generate a Voronoi cell for each entity in the map. The second step is to generate several randomly assigned points in the Voronoi cell of reactions catalysed by proteins. The third step consists in using the k-means algorithm to generate the needed number of clusters and assign the cluster’s centroids coordinates as those of the proteins catalysing the reaction in question.

#### Conversion of obtained coordinates into a standard format (SBML)

##### -Saving protein positions in a BiNoM Reaction Format

Following the previous steps, a file in the BiNoM Reaction Format is obtained, containing the name of the proteins as well as their coordinates and sizes. This simple file will then be converted to a standard CellDesigner SBML format to be compatible with the original metabolic network. As CellDesigner allows the manipulation of “aliases” (multiple copies of the same entity); each protein with the same name present multiple times will have an apostrophe attached to its name based on the number of its repetition within the network.

##### -Conversion of BiNoM Reaction Format into a CellDesigner map

Using a custom python script, information stored in the BiNoM Reaction Format is transformed into a XML file following the SBML format. This file will contain each protein names, IDs, alias IDs, coordinates and type. As for now, only the manipulation of simple proteins is available.

##### -Merging of the ReconMap 2.0 and Proteins maps using BiNoM merging function

Once the file containing proteins to add to the metabolic map is obtained, as they both are in the same SBML format, it is possible to merge them by using a function of the BiNoM plugin. This function allows transforming two or more separated maps into one unique map. This final merged map will be transformed into the NaviCell environment using the NaviCell Factory package (https://github.com/sysbio-curie/NaviCell).

Thus, proteins implicated in the catalysis of a reaction can be seen in the vicinity of the corresponding reactions (Figure 4A). It is important to note that in some cases, reactions are regulated by many proteins, for example in the case of protein families, and the resulting configuration of protein nodes can be very dense (Figure 4B). This aspect can be improved by grouping protein families and visualising them together as a single generic entity. However, it is not always relevant to group all protein sharing a similar name by “family”, since different family members might fulfil distinct or even opposite function, leading to a misinterpretation of the omics data in the context of the maps. Therefore, each protein was kept as a unique and independent entity.

**Figure 4.**
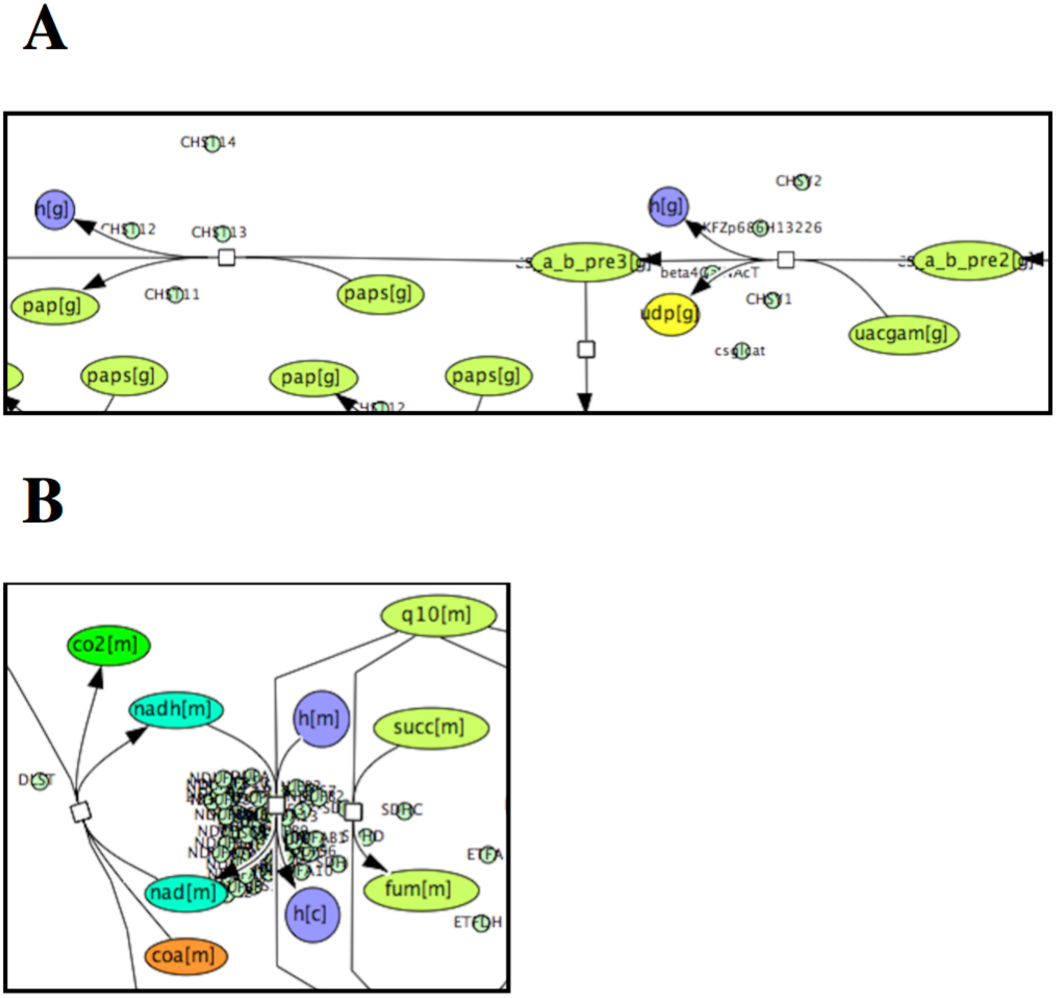
Proteins nodes display on the ReconMap 2.0 canvas. (A) Dispersed distribution of protein nodes around reactions, (B) Case of dense protein nodes distribution around a reaction, implicating 46 proteins. Proteins nodes are represented by pale green circles.

Thanks to this method, 1.550 proteins were allocated in the ReconMap 2.0 canvas associated to more than 7.500 aliases. The algorithm for assigning proteins’ coordinates is robust and its computation time is also scalable as the generation of the 7.500 allocation points is resolved in a matter of seconds.

### 2.4 ACSN-ReconMap 2.0networks integration and visualisation at NaviCell platform

#### ACSN and ReconMap 2.0merging

Once the protein positions file has been generated, it was converted to a CellDesigner [19][20] XML format through a custom python script (https://github.com/sysbio-curie/CellDesigner_networks_map_integration_procedure). This script allows to obtain a file in XML format following the standard of CellDesigner’s SBML. This ‘map’ contains only proteins in the positions they should belong on the final metabolic map. This file was then merged with the ReconMap 2.0 network by using an existing merging function of BiNoM [21][22] to obtain the final network containing the original ReconMap 2.0 as well as the proteins in the vicinity of reactions they catalyse.

#### NaviCell representation of ACNS-ReconMap 2.0 resource

Due to their corresponding size and technical limitations, the two maps couldn’t be merged into one single seamless map. The cross-linking via shared proteins was performed and the two maps were represented as interconnected layers using NaviCell web-based platform, allowing to shuttle between the maps by clicking on a common entity (see next paragraph).

Both maps were preserved with their original layout so that their correspondent relevance of the visual organisation was not lost. Furthermore, this allows users to have an easier view and understanding of the whole system. Moreover, entities annotations from ReconMap 2.0 have been recuperated from the original map and transferring to a NaviCell annotation format (see Materials and Methods), allowing to link entities to corresponding databases (Figure 5).

**Figure 5.**
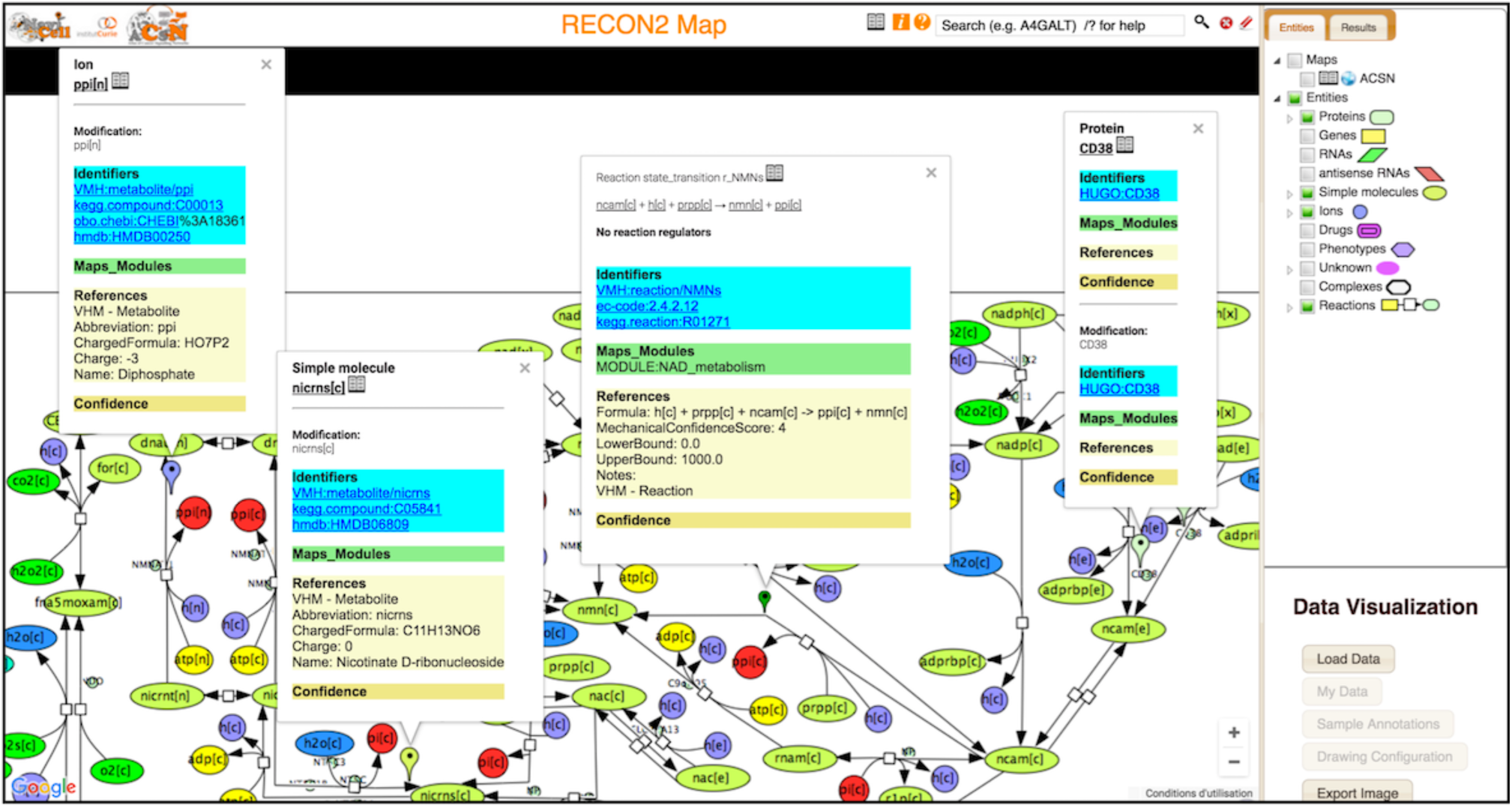
Screenshot of ReconMap 2.0 global metabolic map presented in Google Maps-based interactive environment NaviCell. The map is available at https://navicell.curie.fr/navicell/newtest/maps/ReconMap2/master/index.html.

#### Navigation of ACSN-ReconMap 2.0 resource using NaviCell platform

ACSN and ReconMap 2.0 maps were linked via the common players. Now that proteins had been displayed on top of the ReconMap 2.0 network canvas, those common with ACSN will be used as links to shuttle through both maps. Exploration and shuttling between the two map layers is possible using the NaviCell Google Maps-like features [5]. By clicking on a protein existing in both maps, a window with annotations will appear and a ‘globe’ icon will be clickable in the list seen in the ‘Maps_and_Modules’ section. A new window will be opened and the corresponding protein will be shown on this called map.

In addition, the exploration of the ACSN-ReconMap 2.0 resource is facilitated by the semantic zooming principle of NaviCell platform. As navigating large geographical maps, the semantic zooming on the molecular networks consists in hiding low-level invisible details at a less detailed level of zoom with simultaneous transforming and changing the scale of the representation of the essential objects by creating their abstractions.

This principle can be used for browsing large comprehensive maps of molecular mechanisms such as ACSN [3] and ReconMap 2.0 [1] thanks to the existing open code of Google Maps API.

## 3. Visualisation of cancer multi-omics data in the context of integrated ACSN-ReconMap 2.0 resource

The interconnected ACSN-ReconMap 2.0 resource was applied for visualisation of multi-omic data representing ovarian cancer subtypes. The transcriptomic, copy number and mutation data from TCGA resource was used for the visualisation in the context of the ACSN-ReconMap 2.0 resource using Navicell Web Service toolbox [5], generating molecular portraits of Immunoreactive and Proliferative ovarian cancer subtype. Here below we demonstrate several examples from the molecular portraits and discuss the possible biological significance.

First, we demonstrate that ReconMap 2.0 with displayed protein nodes on the map canvas and provided in NaviCell platform, is now applicable for meaningful multi-omic data visualisation.

As shown in Figure 6, the expression of Keratan metabolism pathway regulators is high in the immunoreactive subtype, where most of the pathway-related genes are mutated and possess a larger copy number variation (Figure 6A). The opposite picture is seen for the Proliferative subtype of ovarian cancer, with mostly under-expressed genes being mutated (Figure 6B).

**Figure 6.**
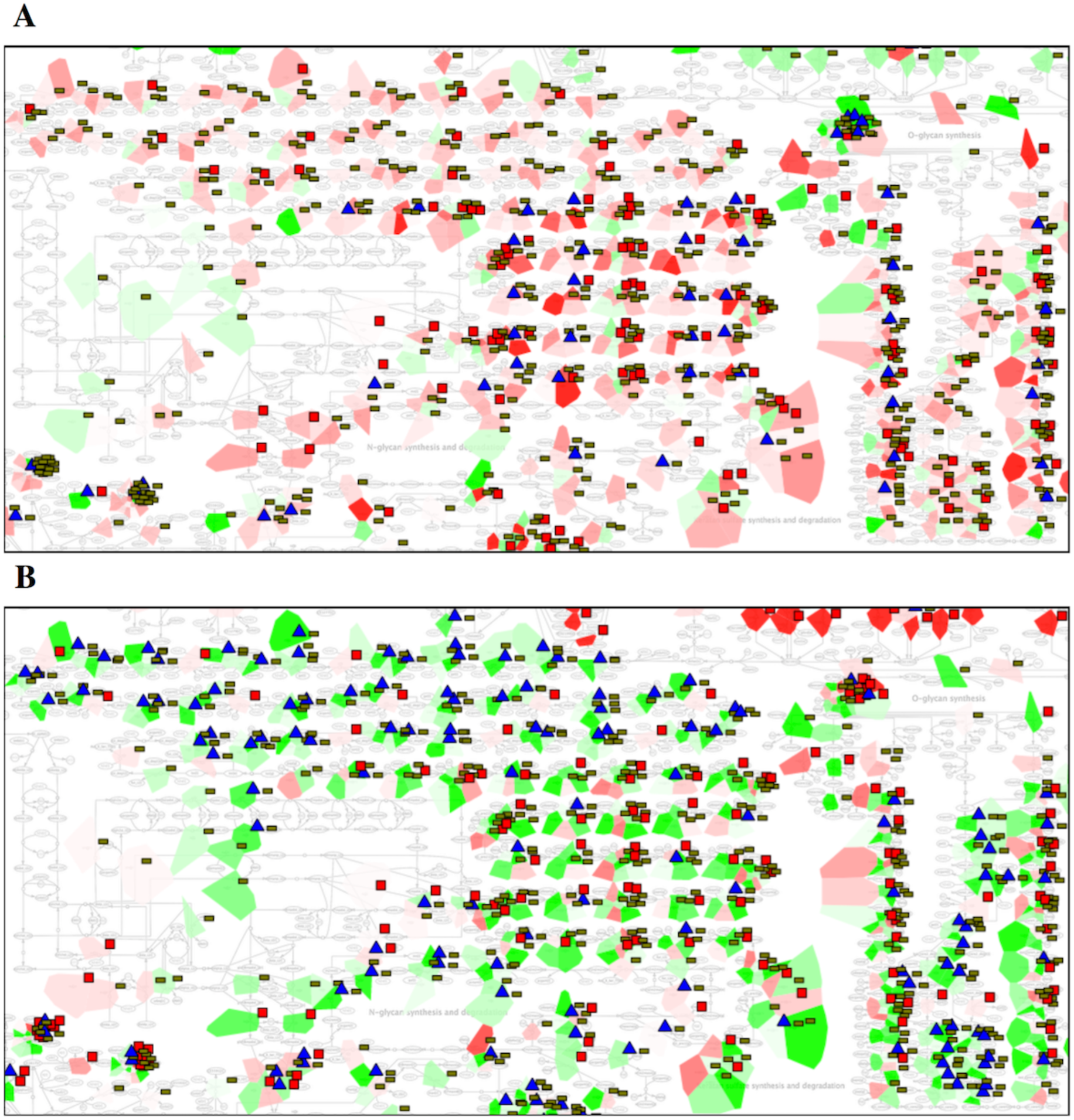
Ovarian cancer multi-omics data visualisation on ReconMap 2.0: zoomed on Keratan sulfate synthesis and degradation metabolic pathway. Two ovarian cancer subtypes are compared: Immunoreactive (top), Proliferative (bottom). Patches using the Map staining function represent the expression level (under-expressed in green and over-expressed in red). Barplots indicate the copy number state (red means at least 2 copy number). Glyphs shown as blue triangles are viewed near genes possessing mutations.

It is known that the Nucleotide metabolism plays an important role in cancer development. The molecular portrait of this metabolic pathway is very different comparing to the Keratan metabolism pathway. The regulators of the Nucleotide transport pathways are under-expressed in the Immunoreactive subtype (Figure 7A) comparing to the in Proliferative subtype (Figure 7B) of ovarian cancer. This observation is in coherence with the fact that highly proliferative cells would consume higher amounts of nucleotides for their growth. Further, for more interpretable visualisation of cross-talks between both maps, only the 252 genes common between ACSN and ReconMap 2.0 were used for visualisations of data.

**Figure 7.**
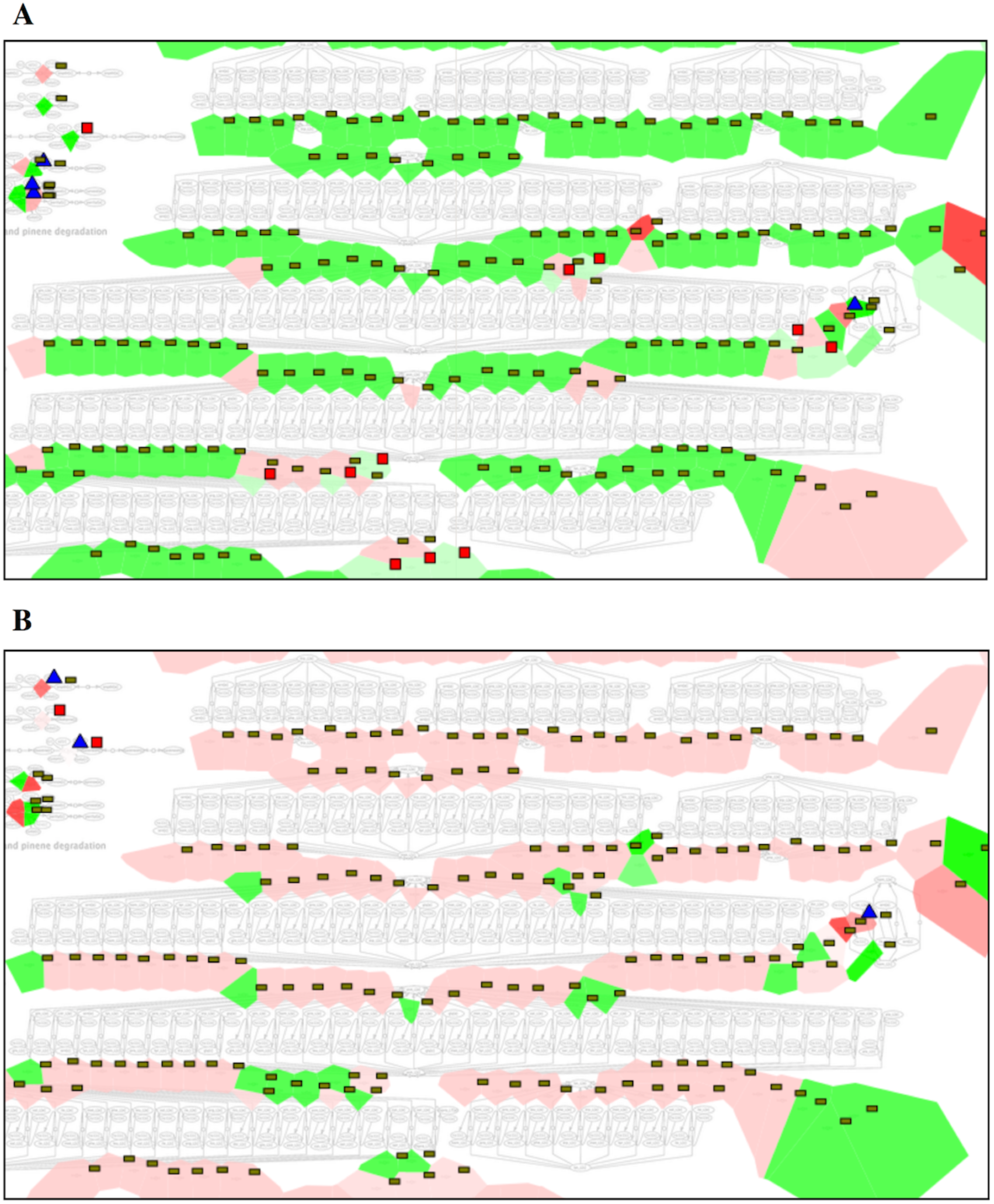
Ovarian cancer multi-omics data visualisation on ReconMap 2.0: zoomed on Nucleotide transport metabolic pathway. Two ovarian cancer subtypes are compared: Immunoreactive (top), Proliferative (bottom) Patches using the Map staining function represent the expression level of the corresponding genes (under-expressed in green and over-expressed in red). Barplots indicate the copy number state (red means at least 2 copy number). Glyphs shown as blue triangles are viewed near genes possessing mutations.

We performed the analysis of the regulation of 252 proteins shared between ReconMap 2.0 and ACSN and retrieved their corresponding implications in the function modules for both maps, comparing two ovarian cancer subtypes as in the previous example. Production of energy is a crucial mechanism necessary for the development of cancer cells, therefore it is not surprising to find significant changes in the regulation of the Energy metabolism module between two ovarian cancer subtypes, especially profound in the Krebs cycle, Glycolysis and Gluconeogenesis mechanisms (Figure 8). In the immunoreactive subtype, genes implicated in the Krebs cycle are over-expressed while those involved in the Glucose metabolism are under-expressed (Figure 8A). The opposite is found in proliferative cells with over-expressed genes in the Glucose metabolism being mutated (Figure 8B).

**Figure 8.**
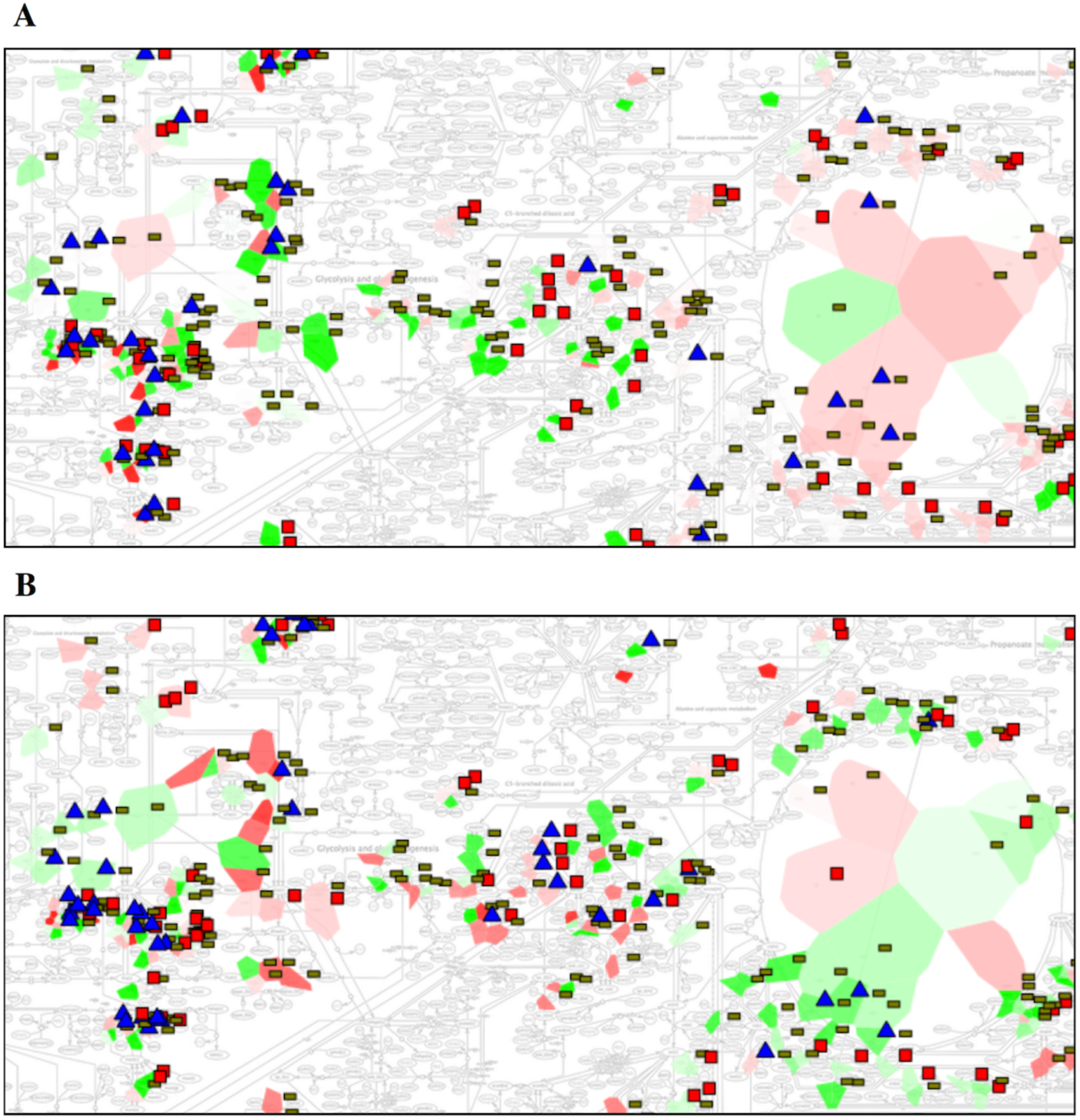
Ovarian cancer multi-omics data visualisation on ReconMap 2.0: zoomed on Energy metabolism pathways. Two subtypes are present: Immunoreactive (top), Proliferative (bottom). Data display modes are as in Figure 5.

The genes implicated in the Inositol phosphate metabolism also show opposite regulation patterns between the two studied groups (Figure 9). This process is known to be dysregulated in cancer and has an impact on cell proliferation and migration [14]. Interpretation of the results in the context of a map can shade light on mechanisms governing these perturbations.

**Figure 9.**
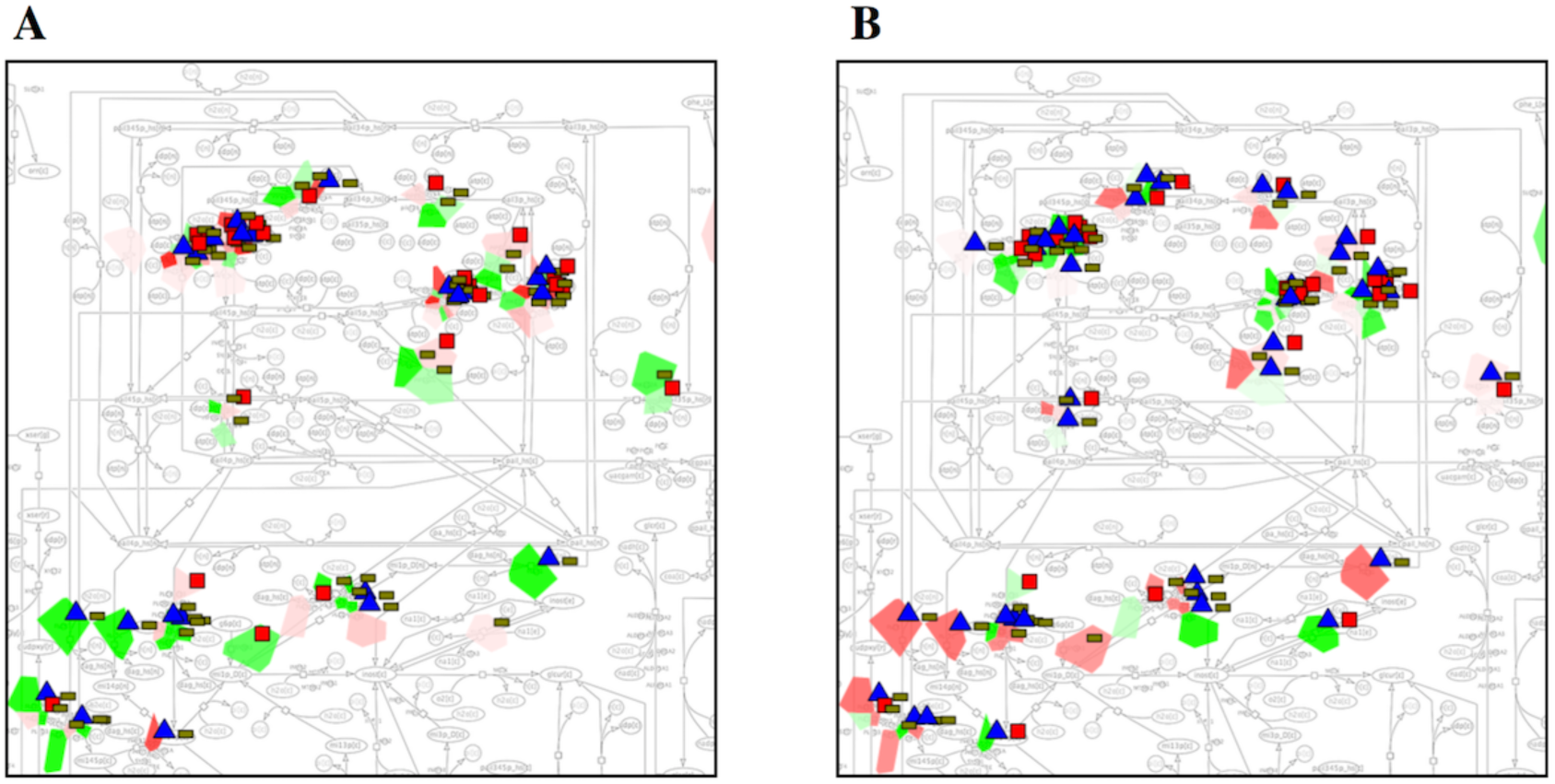
Ovarian cancer multi-omics data visualisation on ReconMap 2.0: zoomed on Inositol Phosphate metabolic pathway. Immunoreactive (A) and Proliferative (B) subtypes are compared. Data display modes are as in Figure 5.

The same type of visualisation has been performed on the ACSN map, demonstrating three deregulated modules: Mitochondrial metabolism (Figure 10), WNT canonical (Figure 11) and WNT no-canonical (Figure 12). These 3 modules are indeed the most enriched in common genes between signalling and metabolic pathways [23].

**Figure 10.**
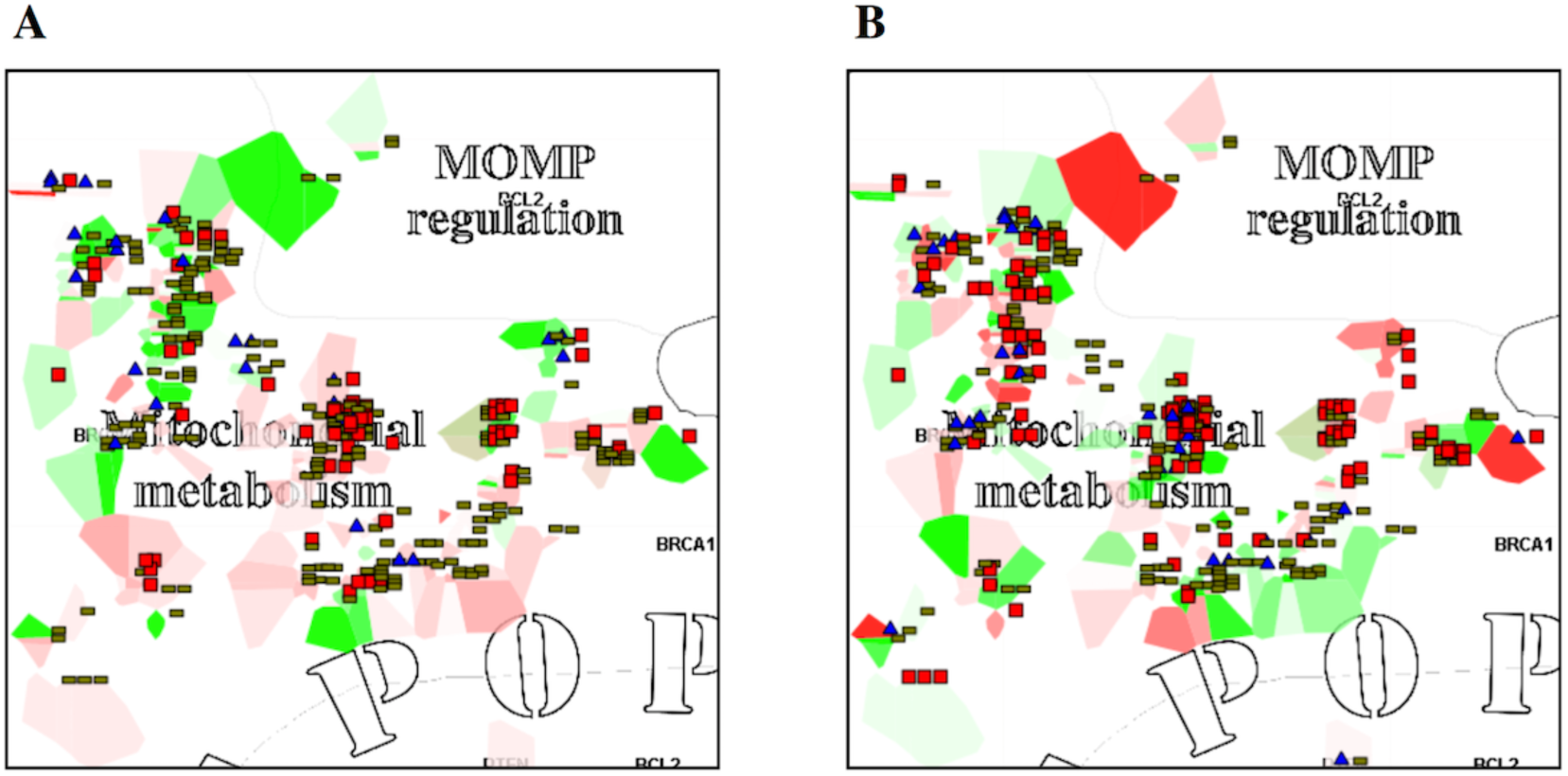
Ovarian cancer multi-omics data visualisation on ACSN: zoomed on Mitochondrial metabolism signalling pathway. Immunoreactive (A) and Proliferative (B) subtypes are compared. Data display modes are as in Figure 5.

**Figure 11.**
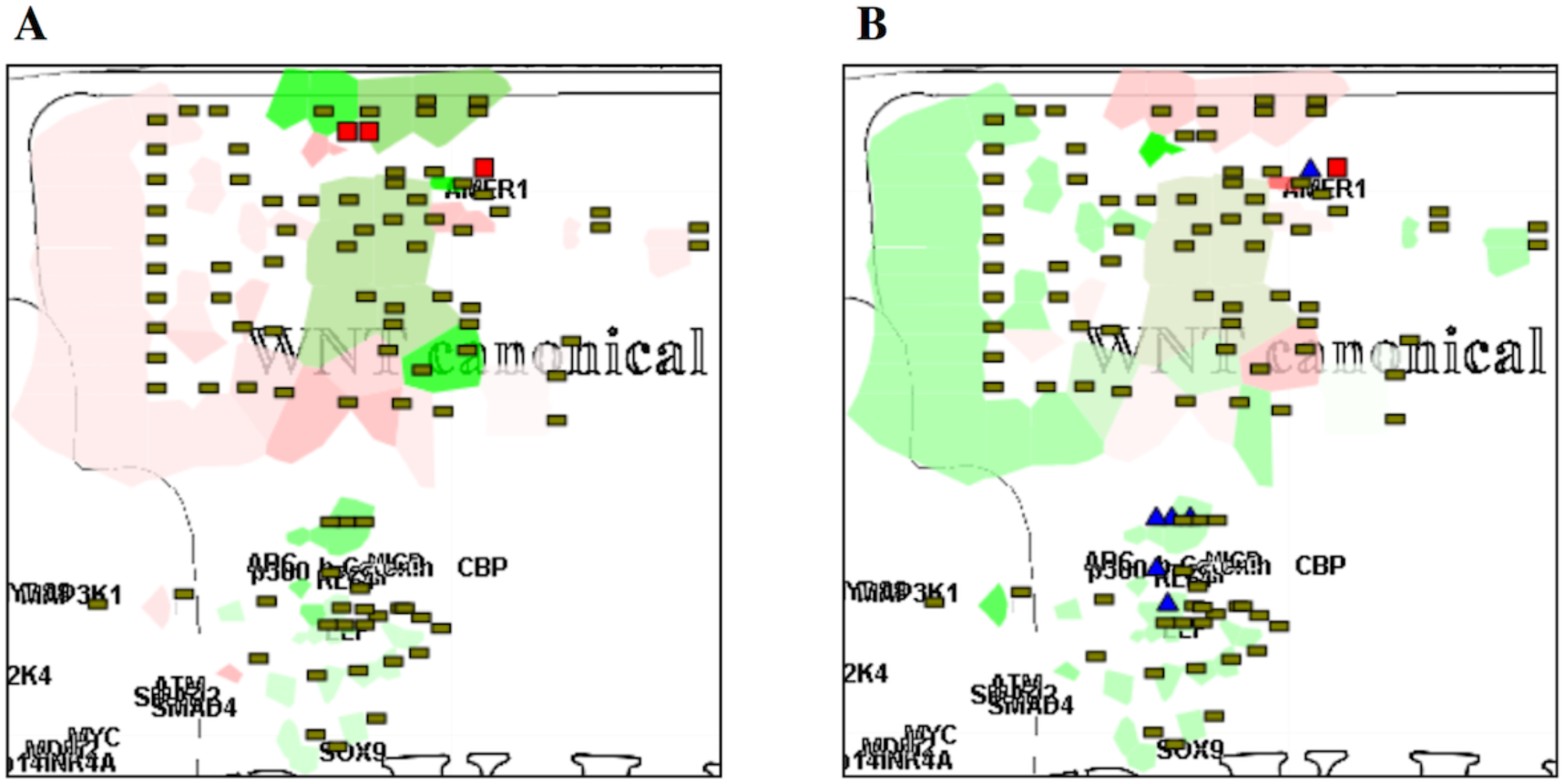
Ovarian cancer multi-omics data visualisation on ACSN: zoomed on WNT canonical signalling pathways. Immunoreactive (A) and Proliferative (B) subtypes are compared. Data display modes are as in Figure 5.

**Figure 12.**
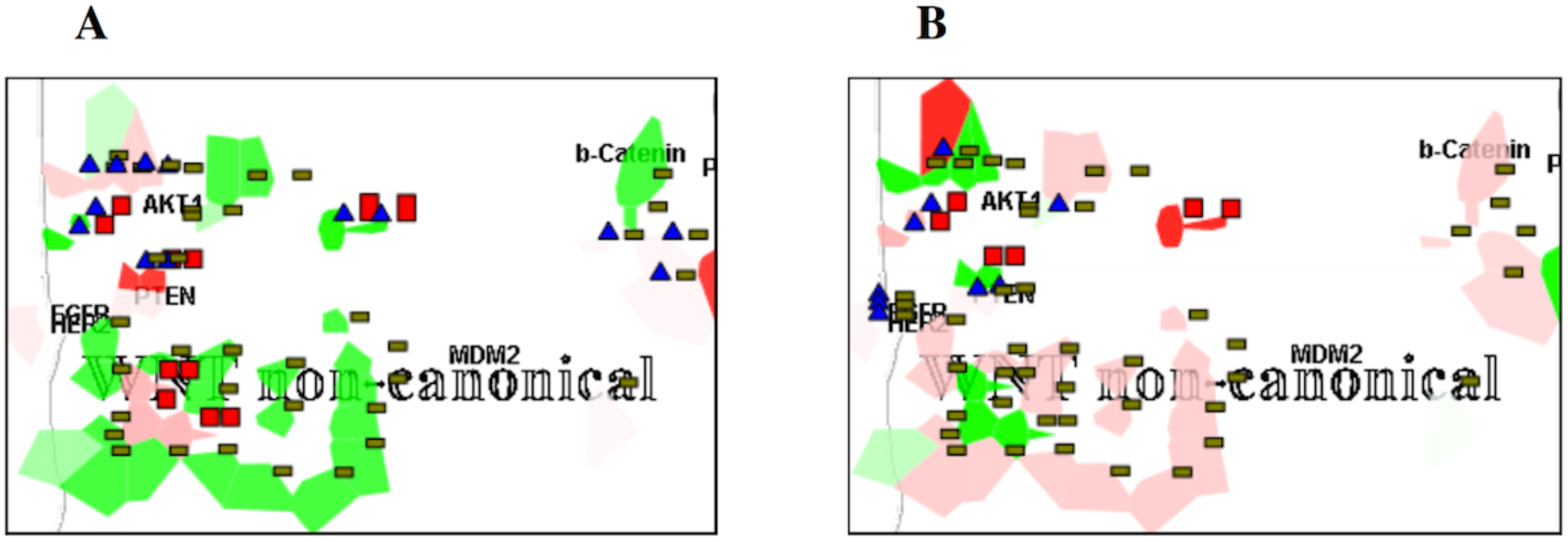
Ovarian cancer multi-omics data visualisation on ACSN: zoomed on WNT non-canonical signalling pathway. Immunoreactive (A) and Proliferative (B) subtypes are compared. Data display modes are as in Figure 5

Interestingly, in WNT canonical and non-canonical modules, the whole cascades seem to be implicated and differ greatly between the two ovarian cancer subtypes. Genes participating in these pathways have been found participating also in the Inositol Phosphate metabolism (Figures 1, 2 and Table 2). This new information is especially valuable because these pathways were not yet shown as related to metabolic processes.

In addition, the data visualisation demonstrated that TNF response factors in ACSN are linked to the Vitamin B2 metabolism in ReconMap 2.0 map and show differential regulation between the two subtypes of ovarian cancer (Figures 1, 2 and Table 2).

## Discussion and conclusions

A systems biology approach involving integration of signalling and metabolic networks permits characterisation of cross-links between the two types of molecular mechanisms in different conditions (e.g. healthy and disease). The integrated ACSN-ReconMap 2.0 resource provided under the NaviCell platform opens an opportunity for a full exploitation of multi-omics data using visualisation features of NaviCell [4]. NaviCell allows to visualise and analyse data based on hierarchical structure of ACSN modules and ReconMap 2.0 subsystems respectively, evaluating ‘activity’ of modules and subsystems thanks to the map staining techniques [5].

ACSN and ReconMap resources are constantly extended with new discoveries in the corresponding fields. Future modifications of these networks will be maintained and the workflow described in this manuscript will be reused to allow updates of the integrated ACSN-ReconMap 2.0 resource.

The developed networks integration methodology and the suggested workflow is a generic mechanism and can be easily applied for integration of other comprehensive maps. The robustness of the method, computational speed and memory usage allows it to be used on any computer with Python and Java installed. Scripts used are open-source and accessible on GitHub (https://github.com/sysbio-curie/CellDesigner_networks_map_integration_procedure).

In this manuscript, we have shown that the merging of metabolic and signalling networks can be achieved and it provides many possibilities for data analyses and comprehension of implicated processes across both maps. In addition, the integrated resource allows to find gaps in connectively between signalling and metabolic processes and suggests exploration of potential links. The integrated ACSN-ReconMap resource will help in further elucidating the crosstalk between of the metabolic and signalling processes and understanding what are the key coordination players in cancer and other human disease.

We will further develop the ACSN-ReconMap resource and integrate into an open software platform together with tools as ROMA [24], COBRA [25], etc. for multi-scale data analysis at morphological, subsystem, reaction and atomic scales. The platform will allow metabolic networks modelling under the regulation of signalling processes with the aim to predict disease status and beyond.

## Methods

### Maps generation tool

CellDesigner [19][20] is a tool used for the construction of both networks and its standard notation allowed the integration and linking across these maps. Both maps are available in a XML format, thus facilitating their automated manipulation.

### Map entity annotation with NaviCell format

The annotation panel followed the NaviCell annotation format of each entity and reaction of the maps includes sections ‘Identifiers’, ‘Maps_Modules’, ‘References’ and ‘Confidence’ as detailed in [3]. ‘Identifiers’ section provides standard identifiers and links to the corresponding entity descriptions in HGNC, UniProt, Entrez, SBO, GeneCards and cross-references in REACTOME, KEGG, Wiki Pathways and other databases. ‘Maps_Modules’ section includes tags of modules in ACSN and metabolic pathways in RecoMap 2, in which the entity is implicated. ‘References’ section contains links to related publications. Each entity annotation is represented as a post with extended information on the entity.

### Generation of NaviCell map with NaviCell factory

NaviCell Factory (https://github.com/sysbio-curie/NaviCell) is a package allowing to convert a CellDesigner map annotated in the NaviCell format into NaviCell Google Maps-based environment. This result in a set of HTML pages with integrated JavaScript code that can be launched in a web browser for online use. HUGO identifiers in the annotation form allow using NaviCell tool for visualisation of omics data [5].

The detailed guidelines for the usage of NaviCell factory, embedded in the BiNoM Cytoscape plugin, is provided at https://navicell.curie.fr/doc/NaviCellMapperAdminGuide.pdf.

### BiNoM

BiNoM (https://binom.curie.fr/) [21][22] is a Cytoscape plugin, developed to facilitate the manipulation of biological networks represented in standard systems biology formats (SBML, SBGN, BioPAX) and to carry out studies on the network structure. BiNoM provides the user with a complete interface for the analysis of biological networks in Cytoscape environment.

### Maps navigation via NaviCell platform

ACSN-ReconMap2.0 interconnected maps are navigable in NaviCell online platform (https://navicell.curie.fr/). NaviCell uses Google Maps and semantic zooming to browse large biological network maps and allows shuttling between the two layers of interconnected resource ACSN-ReconMap 2.0 via common player (proteins).

### Omics visualisation using NaviCell Web Service tool

NaviCell Web Service [5] is a tool for network-based visualisation of ‘omics’ that allows to overlay on maps different types of the molecular data. The tool provides standard heatmaps, barplots and glyphs modes of data display on the maps. In addition, the map staining technique allows to project on the map and grasp large-scale trends in numerical values (such as the whole transcriptome). The web service provides a server mode, which allows automating visualisation tasks and retrieving data from maps via RESTful (standard HTTP) calls.

### Multi-omics data source

The transcriptome, copy number variations and mutation frequencies for ovarian cancer datasets were obtained from the TCGA repository and available at: https://navicell.curie.fr/pages/nav_web_service.html

https://acsn.curie.fr/downloads.html

https://vmh.uni.lu/#downloadview

### Maps accessibility

ReconMap 2.0 in NaviCell format interconnected to ACSN is available at: https://navicell.curie.fr/pages/maps_ReconMap2.html

The ACSN resource is accessible via https://acsn.curie.fr

The ReconMap 2.0 is accessible via https://vmh.uni.lu/#reconmap

## Code accessibility

The code and procedures used for the integration of both networks is accessible on GitHub (https://github.com/sysbio-curie/CellDesigner_networks_map_integration_procedure).

## Acknowledgments

This work has been supported by the COLOSYS grant ANR-15-CMED-0001-04, provided by the Agence Nationale de la Recherche under the frame of ERACoSysMed-1, the ERA-Net for Systems Medicine in clinical research and medical practice and by PRECISE H2020 EU grant. This work received support from MASTODON program by CNRS (project APLIGOOGLE).

## Additional Files

**Additional file 1. List_of_common_proteins.txt.** List of the 252 proteins found in common between ACSN and ReconMap 2.0 maps (available at https://navicell.curie.fr/pages/maps_ReconMap2.html).

**Additional file 2. ACSN_GMT.gmt.** Gene sets composing ACSN modules (available at https://navicell.curie.fr/pages/maps_ReconMap2.html).

**Additional file 3. ReconMap2_GMT.gmt.** Gene sets composing ReconMap 2.0 subsystems (available at https://navicell.curie.fr/pages/maps_ReconMap2.html).

**Additional file 4.** Regulations of reactions in ReconMap 2.0 subsystems by proteins found in ACSN modules. The information encoded in the annotations of reactions and entities on the CellDesigner XML files was analysed and the correspondence between shared proteins in ACSN modules to regulated reactions in ReconMap 2.0 subsystems was retrieved and quantified. Number of ReconMap 2.0 reaction regulated by a subset of shared proteins is shown.

| ACSN Modules | ReconMap 2 metabolic pathways | Number of catalyzed reactions in ReconMap 2 subsystems |
| --- | --- | --- |
| TNF response | Vitamin B2 metabolism<br>Glycerophospholipid metabolism | 3 |
| Caspase | Nucleotide Interconversion | 4 |
| Hedgehog | Inositol phosphate metabolism | 6 |
| MapK | Inositol phosphate metabolism<br>Glycerophospholipid metabolism | 9 |
| PI3K AKT mTOR | Inositol phosphate metabolism<br>Arginine and Proline Metabolism | 13 |
| WNT non canonical | Inositol phosphate metabolism<br>Glycolysis/gluconeogenesis<br>Oxidative phosphorylation | 7 |
| WNT canonical | Inositol phosphate metabolism<br>Nucleotide Interconversion | 18 |
| EMT Motility | Inositol phosphate metabolism<br>Glycerophospholipid metabolism<br>Phosphatidylinositol phosphate metabolism<br>Methionine and cysteine metabolism<br>Selenoamino acid metabolism | 28 |
| Cellcycle | Transport extracellular mainly of Amino Acids<br>Nucleotide Interconversion<br>Folate metabolism<br>Amino Acids metabolism<br>Oxidative phosphorylation<br>Sugar metabolism | 69 |
| Apoptosis genes | Energy metabolism - sugar/amino/citric acid...<br>Extracellular transport of Amino Acids<br>Heme metabolism<br>Inositol phosphate metabolism | 102 |
| Mitochondrial metabolism | Energy metabolism - sugar/amino/citric acid...<br>Extracellular transport in Energy metabolism<br>Fatty acid synthesis<br>Nucleotide Interconversion<br>Amino Acids metabolism<br>Transport mitochondrial<br>Heme metabolism | 295 |

**Additional file 5. Crosstalks_network.txt.** Network of crosstalks based on shared proteins between modules of ACSN and subsystems of ReconMap 2.0 in TXT format. This file contains Source nodes, Target nodes, Interaction type and the number of intersection proteins corresponding to each edge. The file was used for visualisation of the crosstalks in Figure 2 (available at https://navicell.curie.fr/pages/maps_ReconMap2.html).

## References

[1] Noronha A, Daníelsdóttir AD, Gawron P, Jóhannsson F, Jónsdóttir S, Jarlsson S, et al. ReconMap: An interactive visualization of human metabolism. Bioinformatics 2017; 33(4), 605–607.

[2] Thiele I, Swainston N, Fleming RMT, Hoppe A, Sahoo S, Aurich MK, et al. O. A community-driven global reconstruction of human metabolism. Nature Biotechnology 2013; 31(5), 419–425.

[3] Kuperstein I, Bonnet E, Nguyen HA, Cohen D, Viara E, Grieco L, et al. Atlas of Cancer Signalling Network: A systems biology resource for integrative analysis of cancer data with Google Maps. Oncogenesis 2015; 4.

[4] Kuperstein I, Cohen DPA, Pook S, Viara E, Calzone L, Barillot E, Zinovyev A. NaviCell: A web-based environment for navigation, curation and maintenance of large molecular interaction maps. BMC Systems Biology 2013; 7.

[5] Bonnet E, Viara E, Kuperstein I, Calzone L, Cohen DPA, Barillot E, Zinovyev A. NaviCell Web Service for network-based data visualization. Nucleic Acids Research 2015; 43(W1), W560–W565.

[6] Kitano H, Funahashi A, Matsuoka Y, Oda K. Using process diagrams for the graphical representation of biological networks. Nature Biotechnology 2005.

[7] Stoll G, Surdez D, Tirode F, Laud K, Barillot E, Zinovyev A, Delattre O. Systems biology of Ewing sarcoma: A network model of EWS-FLI1 effect on proliferation and apoptosis. Nucleic Acids Research 2013; 41(19), 8853–8871.

[8] Biton A, Bernard-Pierrot I, Lou Y, Krucker C, Chapeaublanc E, Rubio-Pérez C, et al. Independent Component Analysis Uncovers the Landscape of the Bladder Tumor Transcriptome and Reveals Insights into Luminal and Basal Subtypes. Cell Reports 2014; 9(4), 1235–1245.

[9] Jdey W, Thierry S, Russo C, Devun F, Al Abo M, Noguiez-Hellin P, et al. Drug-driven synthetic lethality: Bypassing tumor cell genetics with a combination of AsiDNA and PARP inhibitors. Clinical Cancer Research 2017; 23(4), 1001–1011.

[10] Sassone J, Maraschi A, Sassone F, Silani V, Ciammola A. Defining the role of the Bcl-2 family proteins in Huntington’s disease. Cell Death & Disease 2013; 4(8), e772.

[11] Green and John C, Reed DR, Green DR, Reed JC, Kluck RM, Bossy-wetzel E, Green DR, Newmeyer DD. Mitochondria and Apoptosis. Science 1998; 281(5381), 1309–1312.

[12] Wang C, Youle RJ. The Role of Mitochondria in Apoptosis. Annual Review of Genetics 2009,;43, 95–118.

[13] Tkachuk VA. Phosphoinositide metabolism and Ca2+ oscillation. Biochemistry 1998; 63(1), 38–46.

[14] Tan J, Yu C-Y, Wang Z-H, Chen H-Y, Guan J, Chen Y-X, Fang J-Y. Genetic variants in the inositol phosphate metabolism pathway and risk of different types of cancer. Scientific Reports 2015; 5(1), 8473.

[15] Martinet W, De Meyer GRY, Herman AG, Kockx MM. Amino acid deprivation induces both apoptosis and autophagy in murine C2C12 muscle cells. Biotechnology Letters 2005; 27(16), 1157–1163.

[16] Fu YM, Yu ZX, Lin H, Fu X, Meadows GG. Selective amino acid restriction differentially affects the motility and directionality of DU145 and PC3 prostate cancer cells. Journal of Cellular Physiology 2008; 217(1), 184–193.

[17] Bhargava PM, Allin EP, Montagnier L. Uptake of Amino Acids and Thymidine During the First Cell Cycle of Synchronized Hamster Cells. J. Membrane Biol 1976; (26), 1–17.

[18] Meijer AJ, Dubbelhuis PF. Amino acid signalling and the integration of metabolism. Biochem Biophys Res Commun. 2004; (Vol. 313, pp. 397–403).

[19] Funahashi A, Morohashi M, Kitano H, Tanimura N. CellDesigner: a process diagram editor for gene-regulatory and biochemical networks. BIOSILICO 2003; 1(5), 159–162.

[20] Funahashi A, Matsuoka Y, Jouraku A, Morohashi M, Kikuchi N, Kitano H. CellDesigner 3.5: A versatile modeling tool for biochemical networks. Proceedings of the IEEE 2008; 96(8), 1254–1265.

[21] Zinovyev A, Viara E, Calzone L, Barillot E. BiNoM: A Cytoscape plugin for manipulating and analyzing biological networks. Bioinformatics 2008; 24(6), 876–877.

[22] Bonnet E, Calzone L, Rovera D, Stoll G, Barillot E, Zinovyev A. BiNoM 2.0, a Cytoscape plugin for accessing and analyzing pathways using standard systems biology formats. BMC Systems Biology 2013; 7.

[23] Pate KT, Stringari C, Sprowl-Tanio S, Wang K, TeSlaa T, Hoverter NP, et al. Wnt signaling directs a metabolic program of glycolysis and angiogenesis in colon cancer. The EMBO Journal 2014.

[24] Martignetti L, Calzone L, Bonnet E, Barillot E, Zinovyev A. ROMA: Representation and quantification of module activity from target expression data. Frontiers in Genetics 2016,;7(FEB).

[25] Schellenberger J, Que R, Fleming RMT, Thiele I, Orth JD, Feist AM, et al. Quantitative prediction of cellular metabolism with constraint-based models: The COBRA Toolbox v2.0. Nature Protocols 2011; 6(9), 1290–1307.

